# Effector-independent brain network for auditory-motor integration: fMRI evidence from singing and cello playing

**DOI:** 10.1101/2020.12.24.423508

**Authors:** Melanie Segado, Robert J. Zatorre, Virginia B. Penhune

## Abstract

Many everyday tasks share high-level sensory goals but differ in the movements used to accomplish them. One example of this is musical pitch regulation, where the same notes can be produced using the vocal system or a musical instrument controlled by the hands. Cello playing has previously been shown to rely on brain structures within the singing network for performance of single notes, except in areas related to primary motor control, suggesting that the brain networks for auditory feedback processing and sensorimotor integration may be shared (Segado et al. 2018). However, research has shown that singers and cellists alike can continue singing/playing in tune even in the absence of auditory feedback (Chen et al. 2013, Kleber et al. 2013), so different paradigms are required to test feedback monitoring and control mechanisms. In singing, auditory pitch feedback perturbation paradigms have been used to show that singers engage a network of brain regions including anterior cingulate cortex (ACC), anterior insula (aINS), and intraparietal sulcus (IPS) when compensating for altered pitch feedback, and posterior superior temporal gyrus (pSTG) and supramarginal gyrus (SMG) when ignoring it (Zarate et al. 2005, 2008). To determine whether the brain networks for cello playing and singing directly overlap in these sensory-motor integration areas, in the present study expert cellists were asked to compensate for or ignore introduced pitch perturbations when singing/playing during fMRI scanning. We found that cellists were able to sing/play target tones, and compensate for and ignore introduced feedback perturbations equally well. Brain activity overlapped for singing and playing in IPS and SMG when compensating, and pSTG and dPMC when ignoring; differences between singing/playing across all three conditions were most prominent in M1, centered on the relevant motor effectors (hand, larynx). These findings support the hypothesis that pitch regulation during cello playing relies on structures within the singing network and suggests that differences arise primarily at the level of forward motor control.

**Highlights:**

- Expert cellists were asked to compensate for or ignore introduced pitch perturbations when singing/playing during fMRI scanning.
- Cellists were able to sing/play target tones, and compensate for and ignore introduced feedback perturbations equally well.
- Brain activity overlapped for singing and playing in IPS and SMG when compensating, and pSTG and dPMC when ignoring.
- Differences between singing/playing across were most prominent in M1, centered around the relevant motor effectors (hand, larynx)
- Findings support the hypothesis that pitch regulation during cello playing relies on structures within the singing network with differences arising primarily at the level of forward motor control

## 1 Introduction

Maintaining an intended musical pitch when singing or playing an instrument is a complex sensory-motor skill whose success depends on feed-forward motor control, auditory feedback integration and error correction (Zatorre et al. 2007; Kleber and Zarate 2014). The brain networks underlying auditory-motor integration have been explored in speech and singing (Kleber and Zarate 2014; Rauscheker 2011; Hickok and Poeppel 2004; Tourville and Guenther 2011), but it is not yet known whether the same neural systems underlying vocal control are also engaged for skills like playing instruments, that have common high-level sensory goals to singing, but are phylogenetically newer and depend on very different movements and effectors.

Studies that have separately investigated the brain regions contributing to musical instrument playing and singing often report activation in similar functional networks (Zatorre, Chen, and Penhune 2007; R. M. Brown, Zatorre, and Penhune 2015, Kleber and Zarate 2014). More importantly, a recent fMRI study from our laboratory directly compared singing and cello playing within the same individuals and found that the networks for simple feed-forward control were almost entirely overlapping (Segado et al. 2018). This finding raised the possibility that musical instrument playing may make use of the same auditory-motor networks as vocalization, consistent with the concept of neuronal recycling that posits that culturally newer tasks make use of brain networks that developed for phylogenetically older tasks (Dehaene 2005). However, our previous study examined simple playing and singing, which only engages the feed-forward component of auditory-motor control. A hallmark of auditory-motor integration in the vocal system is feedback control, where auditory feedback is used to modulate the motor response to correct or maintain the appropriate pitch (Burnett et al. 1998). Therefore, the current study again compares singing and cello-playing but uses a pitch perturbation paradigm to test whether similar feedback control and error correction mechanisms are engaged for the two production modalities.

Behaviourally, it has been clearly demonstrated that monitoring of auditory feedback is essential for both singing and playing in tune (J. Chen et al. 2008; Kleber and Zarate 2014). Research on pitch stability in expert cellists during shifting movements has been shown to require ongoing auditory feedback in order to continue producing sequences of notes in tune (J. Chen et al. 2013). Similarly, research on singing has shown that pitch is less accurate and more variable in the absence of auditory feedback (Kleber et al. 2017). However, with sufficient expertise, people can sing and play musical instruments with a high degree of accuracy even when auditory feedback is attenuated (Kleber et al. 2017; J. Chen et al. 2008; Zarate and Zatorre 2008, 2005), likely based on highly developed forward models (Kleber et al. 2013) combined with somatosensory, kinesthetic, and vibrotactile cues (Askenfelt and Jansson 1992; Goebl and Palmer 2008).

Auditory feedback perturbation paradigms, which isolate the effects of auditory feedback on behaviour from those of feedback received through somatosensory modalities, have been used extensively to characterize the role of auditory feedback in auditory-vocal integration. In speech and non-speech vocalizations, it has been shown that participants will reflexively compensate for a variety of auditory feedback perturbations including loudness (Lane and Tranel 1971) and pitch (F0) (Tourville, Reilly, and Guenther 2008). Both of these reflexes are also seen during non-speech vocalization in a variety of non-human animal models as well (J. Luo, Hage, and Moss 2018). A compensatory response to perturbed pitch feedback has also been observed during singing, with research showing that participants compensate for introduced pitch perturbations both when holding a constant pitch (Burnett et al. 1998) and when executing dynamic pitch changes (Burnett and Larson 2002), and that a rapid compensatory response may be involuntary (Zarate, Wood, and Zatorre 2010). Research asking participants to ignore auditory feedback perturbations, thereby requiring them to rely primarily on a forward model of their produced sound based on the remaining sensory feedback (proprioceptive, kinesthetic), has been especially useful in understanding the role of expertise, and has revealed that expert singers are better able to ignore both perturbed pitch (Zarate et al. 2005, 2008) and loudness (Tonkinson 1994) auditory feedback than nonsingers.

Accompanying the behavioural evidence that auditory feedback is used to guide movements for vocal pitch regulation are models that describe the brain networks used for motor planning and execution, auditory feedback processing, and auditory-motor integration (Figure 1). The reciprocal pathway connecting this network of brain regions (auditory, parietal, dorsal pre-motor) comprises the portion of the auditory dorsal stream that we focus on in the present study. The feedforward component includes primary motor, dorsal pre-motor, and supplementary motor cortices (M1, dPMC, SMA) and parts of the cerebellum related to the orofacial articulators (Zatorre 2007; Kleber and Zarate 2014; Rauscheker 2011; Hickok and Poeppel 2004; Tourville and Guenther 2011). Specific regions within M1 vary based on the effector used to accomplish the task with some areas controlling the orofacial articulators (S. Brown 2008) including the larynx, and others controlling bilateral hand movements (Yousry et al. 1997; Lotze et al. 1999). In the vocal system, the periaqueductal gray (PAG), a brainstem nucleus that directly innervates the larynx, has also been found to regulate the initiation of voluntary vocalizations (Larson 1991; Dujardin and Jurgens 2005; Jurgens and Ploog 1970).

**Figure 1.**
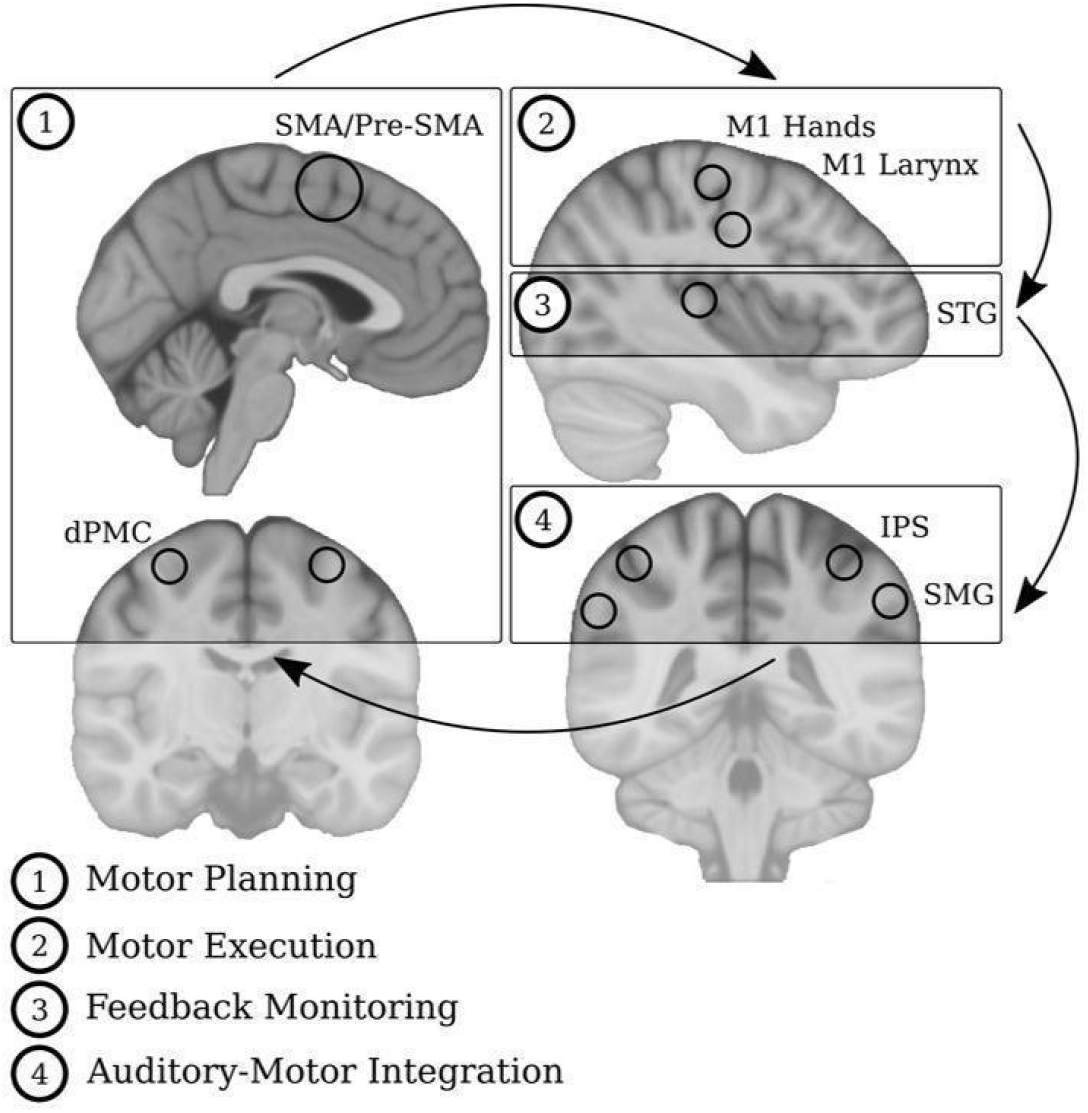
Select components of Feedforward and Feedback networks | Feedforward regions encompass those related to motor planning (SMA/pre-SMA, dPMC), motor execution (M1 Hands, M1 Larynx). Feedback regions encompass those related to feedback monitoring (STG, including HG) and auditory-motor integration (IPS, SMG).

Auditory feedback processing has been shown to involve primary and secondary auditory regions, including Heschl’s gyrus (HG), posterior superior temporal gyrus (pSTG) and superior temporal sulcus (STS). Auditory-motor integration has been linked to pSTG, supramarginal gyrus (SMG), inferior parietal lobule and intraparietal sulcus (IPL, IPS), as well as the anterior insula and anterior cingulate cortex (aINS, ACC) (Zatorre 2007; Kleber and Zarate 2014; Rauscheker 2011; Hickok and Poeppel 2004; Tourville and Guenther 2011). In particular, in a study examining auditory-motor integration during singing, a network involving IPS, SMG, STG/STS, aINS, and ventral pre-motor cortex (vPMC) was found to be engaged when successfully ignoring large pitch feedback perturbations (Zarate et al. 2008), while the ACC and pSTS were found to be engaged during involuntary correction of small pitch shifts (Zarate, Wood, and Zatorre 2010).

The roles of feedforward and feedback brain networks have also been investigated during musical instrument playing. A study on pitch and timing feedback manipulations during keyboard playing found that participants recruited both feedforward (SMA, cerebellum) and feedback (areaSpt, IPL, ACC) areas to a greater extent when pitch feedback was perturbed compared to simply playing a simple melody (Pfordresher et al. 2014). While this study did not allow for the online pitch corrections that occur during continuous pitch instrument performance, this finding still demonstrates that these brain structures are linked to the sound-to-movement transformations necessary for using incorrect pitch feedback to plan future movements. In addition, several studies have found that when participants with no musical training learn to play simple sequences on the keyboard, they quickly form a link between sounds and movements, which is reflected in coordinated activation of motor and sensory brain regions even when only auditory input is provided (J. L. Chen, Zatorre, and Penhune 2006; Stephan, Lega, and Penhune 2018; Lahav et al. 2005; Herholz et al. 2015).

Keyboard studies cannot directly address the role of these structures in pitch regulation since the keyboard is a discrete-pitch instrument, however the same finding has recently been replicated on the cello, which like the voice has a continuous pitch distribution. When novices learned to play a set of simple cello sequences they recruited the auditory to dorsal-cortical pathway after only one week of cello learning compared to the pre-training (Wollman et al. 2018). Moreover, correlated activity between auditory cortex and SMA/pre-SMA after one week of training was found to be predictive of better learning outcomes after 4 weeks of training. The latter finding in particular speaks to the functional importance of sensory-motor links in the production of accurate pitch sequences.

While much is known about the brain networks used for auditory-motor integration, especially for vocalization, the functional role of some brain structures in pitch regulation has not been fully characterized. One such structure is the pSTG. It has been shown that pSTG activity is attenuated in anticipation of self-generated sounds (Bendixen, SanMiguel and Shroger 2012; Mathias, Gehring, and Palmer 2019; Christoffels, Formisano, and Schiller 2007), however studies have also shown that activity in auditory cortex is stronger when listening to sounds produced on the instruments for which one has the greatest expertise (Gebel et al., 2013), and is stronger for expert singers than for non-singers (Zarate et al. 2008), suggesting that expertise may result in enhanced auditory processing in this area (Reznik et al. 2014). Another brain structure whose role is likely critical in sensory-motor mapping is the IPS, which is recruited for high-level pitch transformations such as those required for melody transposition even in the absence of a motor performance task (Foster 2010; Albouy et al. 2017). The IPS has also been shown to be somatotopically organized, such that posterior portions are more involved in visuomotor tasks while anterior portions close to the post-central gyrus are more involved in grasping (Culham and Valyear 2006, Grefkes and Fink 2005).

In the present study we directly investigate whether singing and cello playing use similar or distinct networks of brain regions for pitch regulation. To do this, trained cellists performed a pitch-feedback perturbation paradigm designed to test both the feedforward component of the auditory-motor integration network (ignoring pitch perturbations), and the feedback component (compensating for pitch perturbations). All participants performed both tasks to allow for direct comparisons between cello playing and singing within the same individuals. For the cello playing task, cellists played a fully MR compatible cello device (Hollinger and Wanderley 2013, 2015) used in our previous research (Wollman et al. 2018).

This paradigm allows us to answer specific questions about the role of key brain structures within the singing network. For the IPS, if singing and cello playing overlap directly when compensating for pitch perturbations, then it would support the hypothesis that at least one role of the IPS in pitch regulation is to accomplish high-level pitch transformations, whereas if they are completely separate it would support the hypothesis that its role is to carry out sensory-to-motor transformations specific to the effector needed to accomplish the task, in line with its somatotopy. For the pSTG, contrasting the activity for these two production modalities will allow us to test the hypothesis that cello expertise results in enhanced processing for cello playing relative to singing, or that activity is attenuated for both cello playing and singing due to the anticipation of a self-generated sound. Computing the task-based functional connectivity between an HG seed and areas within the dorsal stream for both singing and cello playing will allow us to test the hypothesis that a tighter coupling between activity in auditory and motor brain regions contributes to performance. More generally, the present study will allow us to extend existing theories about cortical representation of complex skills, including the theory of neuronal recycling, by testing whether pitch regulation during instrument playing, a human-specific cultural task, uses phylogenetically ancient brain networks that exist for vocal pitch regulation.

## 2 Materials and methods

### 2.1 Subjects

A total of 15 expert cellists (10 female) were recruited from the Montreal community. They reported normal hearing, no neurological disorders, and had no contraindications for the MRI environment. Thirteen participants (9 female) were included in the final analysis (mean years experience = 12.93 ± 4.46, mean starting age = 9, mean practice hours per week = 12.07); two were excluded from the analysis due to technical problems. 11 of 13 participants included in the analysis were enrolled in graduate or undergraduate cello performance programs that included solfege and choral training. However, potential participants that reported formal vocal training beyond the required courses were not selected to participate. The remaining two participants had more than 10 years cello experience, but were not enrolled in a university program. This study was approved by and carried out in accordance with the recommendations of Montreal Neurological Institute Research Ethics Board and the McConnell Brain Imaging Centre. All subjects gave written informed consent in accordance with the Declaration of Helsinki.

### 2.2 Experimental Paradigm

#### 2.2.1 Task Conditions

There were three main task conditions (Compensate, Ignore, and Simple) for each of two production modalities (Cello, Singing). In each of the three conditions, participants were presented with a 1 s auditory target tone (two target tones were used throughout the trials) that they were instructed to sing or play back for 2.5 s (Figure 2A). Participants heard both the auditory targets and the auditory feedback of their own performance binaurally through insert earphones. A pink-noise background was used to mask bone conduction so that participants could only use the auditory information provided through the earphones to the greatest extent possible. For Compensate and Ignore trials (40% of trials, 240 total), between 1 and 1.5 s after participants started singing/playing back the target, the pitch of their auditory feedback was shifted up or down by 100 cents (one semitone) in an unpredictable manner (jittered up to 500 ms). On Compensate trials, participants were instructed to make a compensatory movement in order to go back to hearing themselves sing/play the intended (target) tone (Figure 1B). On Ignore trials, participants were instructed to avoid making any compensatory movements, and to continue singing/playing the target tone even though its pitch would sound shifted in their earphones (Figure 2B). The Simple condition comprised the subset of trials within the Compensate or Ignore conditions where no shift was introduced (20% of trials, 120 total) and thus participants simply had to sing or play back the presented target tone without perturbation.

**Figure 2.**
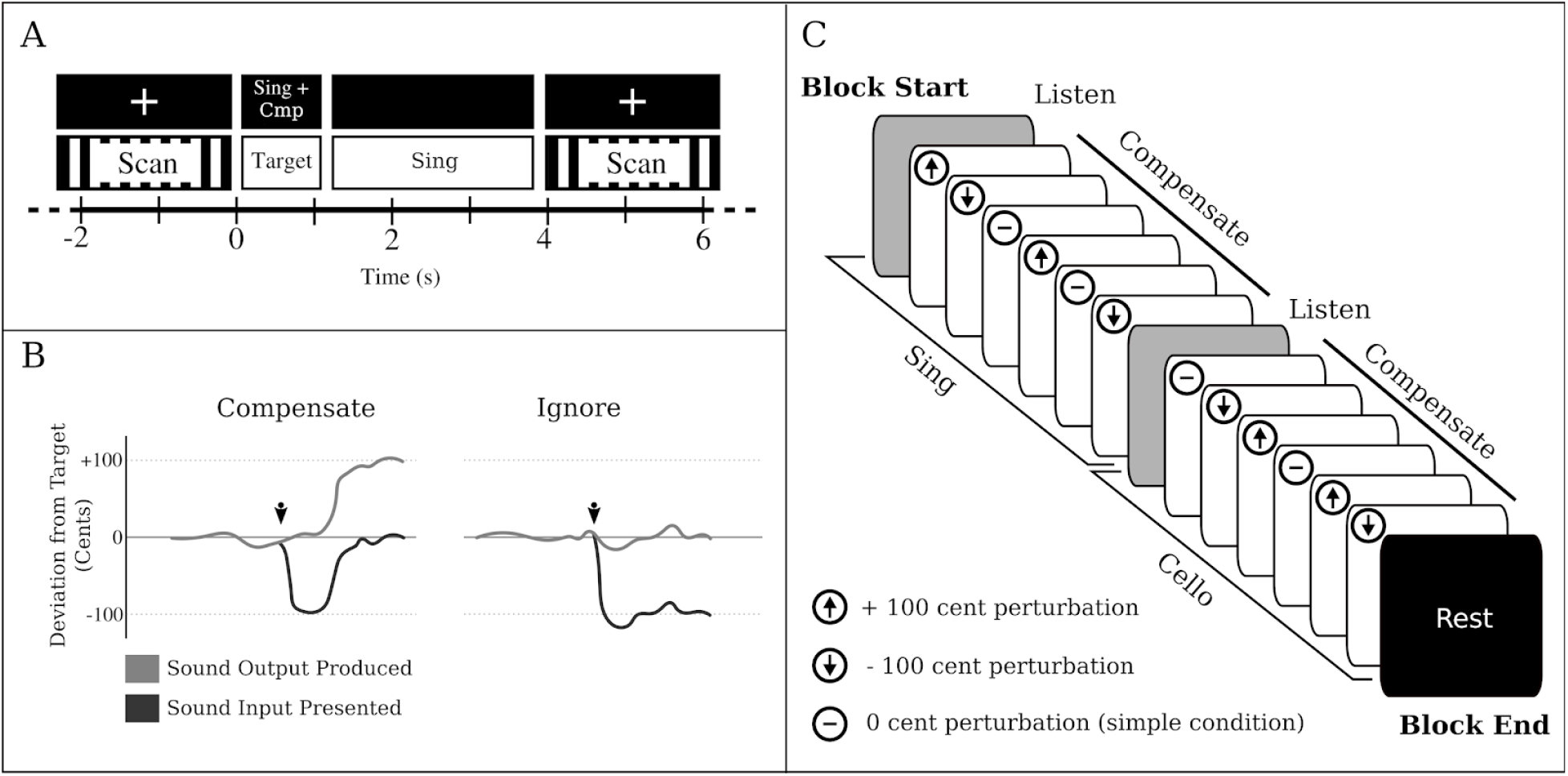
Task and Conditions | A) Example of one experimental trial. Sparse sampling design used to avoid auditory and motion artifacts. Auditory target presentation, instruction presentation, and singing and cello playing were done during the silent period between scans B) Typical behavioural response to pitch feedback perturbation when participants were instructed to Compensate (left) or to Ignore (right) for an introduced pitch perturbation of 100 cents (arrow indicates shift onset) C) Example of one 15-trial Compensate block. Instrument and Condition were counterbalanced across runs, and note/pitch perturbations were pseudorandomized within each block. Simple condition trials (perturbation magnitude 0 cents) were interspersed within Compensate/Ignore blocks.

Three additional control conditions were included in this experiment: Masked Feedback, Listen, and Rest. In the Masked Feedback condition (20% of trials, 120 total), participants were presented with the same target tones as in the Simple/Compensate/Ignore conditions and were asked to sing/play them back but they received only pink masking noise through the earphones. For Listen trials (13.33% of trials, 80 total) participants were visually cued to listen, and the auditory target tone was followed by a 2 s pre-recorded, non-pitch shifted playback of that target. Participants were instructed to listen to the playback of the tone without making any movements or sounds. For Rest trials (6.67% of trials, 40 total) participants were visually cued to rest, and were instructed to not move or make any sounds.

#### 2.2.2 Stimuli

Presented target tones corresponded to pitches from the Western music scale E3, F#3, for cello, and E3, F#3, or E4, F#4 (one octave higher) for singing depending on the participant’s vocal range. Tones were recorded by either a female vocalist, male vocalist, or on a cello. Female participants heard and imitated female voices, male participants heard and imitated male voices. At the start of each session, sound levels were set to default values of 78.3db SPL for pink masking noise and 15.6 dB above the noise floor for auditory targets. Volume settings were adjusted on a per-subject basis such that masking noise and auditory targets were presented at a comfortable volume and that masking noise attenuated non-pitch shifted feedback to the greatest extent possible. To determine this volume level, a pitch shift of 100 cents was applied and participants would sing/play a constant note while the experimenter adjusted the volume until participants reported only hearing the pitch shifted note through their earphones. The position of volume knobs was noted for the subsequent fMRI session.

### 2.3 Experimental Design

The experiment comprised 10 fMRI runs, each of which had 4 blocks of 15 trials. Each block was made up of a single condition (Compensate, Ignore, Masked Feedback) divided into sets of 6 trials on each production modality (cello, singing). Each set of 6 trials was preceded by 1 Listen trial, and a single Rest trial was included at the end of each block (Figure 2C). The order of instructions and production modalities was counterbalanced across runs, and the pitch perturbation (2 no-shift, 2-shift up, 2-shift down) was pseudo-randomized within Compensate/Ignore blocks.

A sparse sampling paradigm was used for the fMRI session (Belin et al. 1999), where a long delay in TR was used to allow tasks to be carried out in the silent period between functional volume acquisitions, thus minimizing acoustical interference. Sparse sampling can also reduce movement-related artifacts since the scanning takes place after the motor production for each trial. The start time of individual trials was jittered up to 500 ms to increase the likelihood of catching the peak of the haemodynamic response.

#### 2.3.1 Procedure

To allow participants to adjust to the fMRI-compatible cello and the constraints of playing it in the scanner, each person underwent a 10 min familiarization session no more than 1 week prior to their session in the MR scanner. During the familiarization session each participant was asked to lie inside a structure that simulated the space constraints of the MRI environment. For both the familiarization session and the fMRI session, a microphone (Optimic 2150, Optoacoustics) was suspended approximately 5 cm from the mouth, the MRI compatible cello device was placed along the torso using a specialized support, and bilateral sound delivery was provided via insert earphones (Sensimetrics, Dayton Audio DTA-1 amplifier). The microphone and cello were connected to a Zoom F8 portable field recorder, and to a midi controlled TCHelicon VoiceOne pitch shifter. The pitch shifter, which was controlled via a USB to midi cable from the experiment control computer, was used to shift the pitch feedback on some trials, or mask the feedback on other trials (Figure 3). All participants were asked to play both target notes on the cello and to sing both target notes, and to try warming up as they would on a regular instrument. Participants also had ample practice on the experimental task prior to the fMRI session. During the familiarization session, participants performed one practice block each of the compensate and ignore conditions. In addition, as part of another study, participants were all trained on and performed 30 blocks of the compensate and ignore tasks in the week prior to the fMRI session.

**Figure 3.**
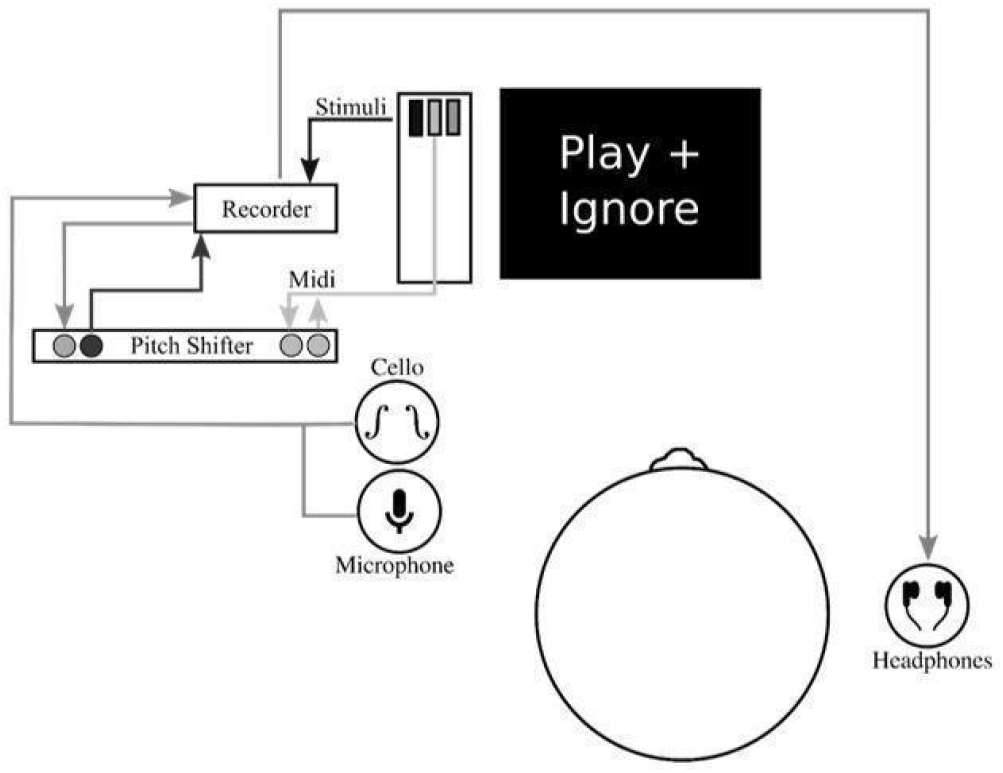
Experimental Setup | Experimental setup used to present stimuli, cello audio, and singing audio through headphones. Experiment control computer displayed visual cues via DVI connector, presented auditory stimuli (target tones) and masking noise through audio jack, and controlled pitch shifter parameters via USB-Midi cable. The pitch shifter, which received inputs from the cello and mic via the portable recorder, allowed for audio feedback from cello/singing to be shifted or attenuated on specific trials while still presenting audio stimuli and masking noise. All audio channels (shifted audio + stimuli + masking noise) were mixed on the portable recorder and presented through headphones after amplification.

Within 1 week of the familiarization session, participants were tested in the Siemens Trio 3T magnetic resonance (MR) scanner at the Brain Imaging Center of the Montreal Neurological Institute. Each participant was fitted with MR-compatible earphones. The MR-compatible microphone was attached to the mirror support system and the MR-compatible cello was laid across the torso using a special MR-compatible stand that prevented it from moving during task performance.

#### 2.3.2 MRI Acquisition

During the 10 functional runs, one whole-head frame of 28 contiguous T2*-weighted images were acquired (Slice order = Interleaved, TE = 85 ms, TR = 6.7 s, Delay in TR = 4.4 s, 64 × 64 matrix, voxel size = 4mm^3^). All tasks were performed during the 4.4s silent period between functional volume acquisitions. On each trial, the target tone was delivered for one second together with the visual instruction; the rest of the trial was taken up by reproduction of the cello or sung tone, or by no action on Rest trials (See Figure 2B). A high-resolution whole-brain T1-weighted anatomical scan (voxel size = 1mm^3^) was collected between runs 5 and 6.

#### 2.3.3 Behavioral Analyses

Individual trials of singing and cello playing were analyzed using the pitch information extracted from the recorded audio signals. Each trial’s audio was first segmented from the continuous 6-track audio using Audacity software. Recorded audio was used to rule out the possibility that participants were singing or humming when instructed to play cello, or playing the cello when instructed to sing. The trials were then processed using a custom analysis pipeline implemented in Python, with a GUI for visualizing and optimizing analysis parameters.

The ambient noise in the scanner room had a peak resonance of 160 Hz that interfered with the extraction of fundamental target pitches, so harmonics 3–10 of the cello and singing tones were used for pitch extraction. To reject room noise and to isolate the harmonics of interest, the raw microphone signal was high-pass filtered with a cutoff at 367 Hz and low-pass filtered with a cutoff at 4,216 Hz. Pitch estimation was then performed using the YinFFT algorithm provided in the Python module Aubio (Brossier 2007), producing a time-series of pitch estimations (detected harmonic, in Hz) and confidence ratings (between 0 and 1). Estimates were adjusted to their representative fundamental pitches before selecting stable pitch regions for further analysis. Stable pitch regions were defined as: segments of at least 150 ms in which the rate of pitch change did not exceed 100 Hz/s (or approximately 0.07 Hz per 32-sample pitch estimation window at the sampling rate of 44,100 Hz). Of these regions, only those that maintained a confidence rating of at least 0.7 according to the algorithm were included. Trials were rejected if no regions were found to meet the stability and confidence criteria, if participants started playing after the pitch shift was introduced, or if they stopped playing before the end of the trial. In total, 86.5% of trials were retained. Rejected trials were excluded from the fMRI analysis.

Pitch extraction was performed for both the unshifted raw output track (produced) and for the shifted headphone track (perceived) to allow for a direct comparison between the produced and perceived audio. Our main outcome measure, deviation from the expected tone, was calculated as the difference between the produced tone and the target tone (expressed in cents, or 100^ths^ of a semitone) within three 150 ms windows relative to shift onset (0 ms): Pre-Shift (− 150 ms to 0 ms), Early Post-Shift (150 ms to 300 ms), and Late Post-Shift (1000 ms to 1150 ms) for each of the Simple, Compensate, and Ignore conditions (Equation 1). For the simple condition, where no shift was introduced, timepoints were chosen relative to the midpoint of the production audio within the trial. Because we had no specific hypotheses related to the directionality of these perturbations, upwards and downwards perturbations were treated as equivalent, and values from the downwards perturbation were therefore multiplied by −1.

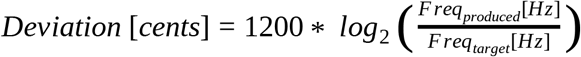

Equation 1 Conversion from frequency ratio (in hertz) to deviation from the target tone expressed in cents (cent deviation) where 100 cents = one semitone.

In addition to the trial-by-trial pitch traces, a global measure of accuracy for each participant was calculated to examine the relationship between overall performance and brain activity. This measure was determined by averaging the cent deviation from the target tone at the Pre-Shift and Late Post-Shift timepoints, including the Simple condition when no pitch shift was introduced so as to account for any natural drift in pitch throughout the trial. For the Compensate condition, perfect performance was considered to be 100 cents in the direction opposite to the introduced pitch perturbation, whereas for the Ignore condition perfect performance was considered to be 0 cents deviation from the target tone. In order to examine BOLD activity as a function of task performance, scores across trials for each condition were averaged to give a global participant score, which was used in the regression analyses.

#### 2.3.4 fMRI Analyses

fMRI data were analysed using the FSL6.0 FEAT toolbox (Jenkinson et al. 2012). Brain extraction was carried out using BET2. Functional volumes were aligned to the high resolution anatomical and then to MNI152 1mm standard space using FLIRT non linear registration with 12 degrees of freedom and a warp resolution of 10mm. To improve the registration quality, b0 unwarping was carried out using a percent signal loss threshold of 50%. Motion parameters were estimated using MCFLIRT, and fMRI time course was temporally filtered to remove drifts greater than 160 ms. To further decrease the number of motion-related active voxels outside the brain, a brain mask based on the MNI152 1mm brain was applied prior to thresholding. To boost the signal to noise ratio, images were spatially smoothed with an 8 mm FWHM kernel. FLAME-1 mixed effects modeling was used to fit the GLM to the fMRI signal.

Statistical significance was determined using an FSL cluster probability threshold of *p* < 0.05 with a voxel-wise significance level of *z* = 2.3 (*p* < 0.05). The cluster probability threshold serves as a correction for multiple comparisons. A total of 21 contrasts were carried out. Contrasts were carried out for each of the instruction conditions (simple vs rest, compensate vs simple, and ignore vs simple) within each of the production modalities (singing, cello playing). Additional contrasts were carried out to directly compare the two production modalities within each of the instruction conditions (e.g. singing compensate vs cello compensate), and between instruction conditions ([singing compensate vs ignore] vs [cello compensate vs ignore]). Additionally, task performance (on a per-subject basis) was regressed against the BOLD signal for each of these contrasts to determine whether or not activity in these areas is positively correlated with good performance on the task, as would be expected if the region in question is specifically linked to the cognitive/motor demands of the task of interest. Statistical conjunctions were carried out to identify commonalities in singing vs cello playing, using the conjunction script created by the Warwick University Department of Statistics, which also made use of the FSL tools (Nichols et al. 2005). This script carries out a voxel-wise thresholding of *p* < 0.05 in both conditions of interest, and then carries out a cluster correction of *p* < 0.05.

Functional Connectivity analyses were carried out using the FSL6.0 FEAT toolbox (Jenkinson et al. 2012). A seed region in the region of primary auditory cortex [Heschl’s Gyrus (HG)] was identified by masking the conjunction of singing and cello playing from the functional data with an anatomically defined mask (Harvard Structural Brain Atlas). The activation time course in the HG seed was extracted and correlated with the whole-brain time course for each task of interest, which was estimated using the GLM. Correlated voxels were thresholded as described above. Regions that showed a correlated time course were then correlated with task performance to determine whether those areas were contributing directly to good performance on the task.

## 3 Results

### 3.1 Behavioural Findings

To determine how accurately participants were able to match the target tone, pitch was averaged across all three timepoints within the simple condition and the cent deviation between the average pitch and the target tone was computed (mean = 0.86 ± 0.27 cents). A 3-way repeated measures anova (2 production modalities x 3 condition x 3 timepoints) was carried out to determine the extent to which participants could compensate for introduced pitch perturbations and also the extent to which they could ignore them both when singing and playing the cello. The analysis showed a significant main effect of condition (F(2,24)=256.79, p < 0.001), a significant main effect of timepoint (F(2,24)=278.5, p < 0.001), and a significant condition x timepoint interaction (F(4,48)=151.91, p < 0.001). There was no significant main effect of production modality (F(1,12)=3.83, p > 0.074), no production modality x condition interaction (F(2,24)=0.92, p > 0.41), no production modality x timepoint interaction (F(2,24)=0.42, p > 0.66), and no production modality x condition x timepoint interaction (F(4,48)=0.26, p > 0.9). Pairwise multiple comparisons (Bonferroni corrected) were carried out to look at the simple main effects of timepoint and condition within each production modality (Figure 4).

**Figure 4.**
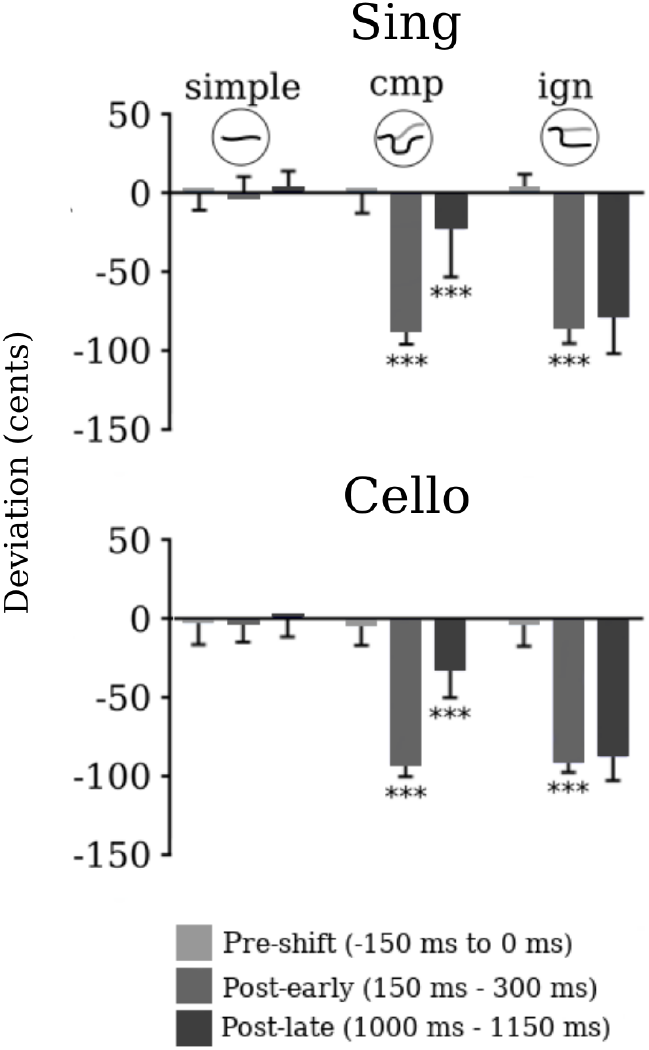
Deviation from target tone during simple, compensate, and ignore conditions | Deviation from target tones at three 150 ms windows (Pre-shift, Post-early, Post-late) relative to pitch perturbation onset (0 ms) for singing (top) and cello playing (bottom) as heard through the participant’s headphones. In simple, cmp, and ign circles, black trace represents what participants heard through headphones and grey trace represents what participants produced. For the Simple condition where no shift was introduced the ‘shift’ was considered to be at the midpoint of the produced audio. For Compensate and Ignore conditions, perturbation was 100 cents. Significance stars refer to cent change from the previous time point.

#### 3.1.1 Singing

Within the singing production modality (Figure 4 top subplot), there were no significant differences across conditions at the pre-shift timepoint [pre_simple_ = pre_ign_ (p > 0.062, 95% CI of the difference = −7.57 to 0.16), pre_simple_ = pre_cmp_ (p > 0.99, 95% CI of the difference = −4.33 to 4.1), pre_cmp_ = pre_ign_ (p > 0.42, 95% CI of the difference = −9.9 to 2.72)], demonstrating that participants were all able to match the target tone before the manipulation was introduced (mean_pre_=0.86 ± 18.09 cents). Across conditions for the early post-shift timepoint, analyses showed a significant difference for simple vs ignore (post-early_simple_ > post-early_ign_, p < 0.001, 95% CI of the difference = 72.64 to 99.29) and simple vs compensate (post-early_simple_ > post-early_cmp_, p < 0.001, 95% CI of the difference = 75.41 to 100.98), but no significant difference for compensate vs ignore (post-early_cmp_ = post-early_ign_, p > 0.29, 95% CI of the difference = −5.68 to 1.22). This indicates that 150 ms following the introduced pitch shift, participants were equally affected by the pitch change in the compensate condition (mean_post-early(cmp)_=-87.51 ± 14.86) as in the ignore condition (mean_post-early(ign)_=-88.86 ± 12.99).

Within the simple condition, no significant differences were observed for the comparison of pre-shift vs post-early (pre_simple_ = post-early_simple_, p > 0.79, 95% CI of the difference = −2.72 to 1.1) or post-early vs post-late timepoints (post-early_simple_ = post-late_simple_, p > 0.46, 95% CI of the difference = −9.93 to 2.89) showing that participants were able to sing a stable pitch when no feedback manipulation was introduced (mean_simple_ = −1.09 ± 19.42 cents). For the compensate condition participants showed a significant pitch deviation from pre-shift to the early post-shift timepoint (pre_cmp_ > post-early_cmp_, p < 0.001, 95% CI of the difference = 72.85 to 102.16) and then a significant reduction in deviation from the early to the late post-shift timpoint (post-early_cmp_ < post-late_cmp_, p < 0.001, 95% CI of the difference = −91.42 to −39.21), indicating that they both responded to and were able to compensate for the perturbation. For the ignore condition, participants showed significant pitch deviations for the early post-shift timepoint (pre_ign_ > post-early_ign_, p < 0.001, 95% CI of the difference = 77.45 to 100.27) that did not change for late post-shift timepoint (post-early_ign_ = post-late_ign_, p > 0.67, 95% CI of the difference = −21.19 to 7.8). The compensate vs ignore comparison was significant in the post-late timepoint (post-late_cmp_ > post-late_ign_, p < 0.001, 95% CI of the difference = 28.97 to 83.81) indicating that participants responded to the perturbation and were able to maintain a stable pitch in the face of conflicting feedback when instructed to ignore.

#### 3.1.2 Cello Playing

The same set of analyses carried out for singing were also performed for cello playing (Figure 4 bottom subplot) and the results paralleled those observed for singing. The pairwise contrasts showed no significant differences across conditions at the pre-shift timepoint [pre_simple_ = pre_ign_ (p > 0.99, 95% CI of the difference = −2.86 to 4.95), pre_simple_ = pre_cmp_ (p > 0.99, 95% CI of the difference = −3.91 to 7.09), pre_cmp_ = pre_ign_ (p > 0.99, 95% CI of the difference = −6.19 to 5.1)], demonstrating that participants were all able to match the target tone before the manipulation was introduced (mean_pre_=-3.53 ± 23.52 cents). Across conditions for the early post-shift timepoint, analyses showed a significant difference for simple vs ignore (post-early_simple_ > post-early_ign_, p < 0.001, 95% CI of the difference = 77.96 to 98.84) and simple vs compensate (post-early_simple_ > post-early_cmp_, p < 0.001, 95% CI of the difference = 78.26 to 101.34), but no significant difference for compensate vs ignore (post-early_cmp_ = post-early_ign_, p > 0.99, 95% CI of the difference = −7.14 to 4.34). This indicates that 150 ms following the introduced pitch shift, participants were equally affected by the pitch change in the compensate condition (mean_post-early(cmp)_=-88.19 ± 14.92) as in the ignore condition (mean_post-early(ign)_=-87.34 ± 15.76).

Within the simple condition, pitch was stable across all three timepoints with no significant differences observed for the comparison of pre-shift vs post-early (pre_simple_ = post-early_simple_, p > 0.99, 95% CI of the difference = −2.15to 2.12) or post-early vs post-late timepoints (post-early_simple_ = post-late_simple_, p > 0.99, 95% CI of the difference = −6.3 to 1.48) (mean_simple_ = −1.83 ± 21.58 cents). Within the compensate condition, significant differences were observed for pre-shift vs post-early (pre_cmp_ > post-early_cmp_, p < 0.001, 95% CI of the difference = 76.68 to 99.7) and post-early vs post-late timepoints (post-early_cmp_ < post-late_cmp_, p < 0.001, 95% CI of the difference = −72.61 to −47.23). As when singing, participants compensated for the feedback manipulation (mean_cmp_ = −59.92 ± 19.28 cents) in the compensate condition, but not in the ignore condition (mean_ign_ = −3.77 ± 16.84 cents). There was a significant difference between pre vs post-early timepoints (pre_ign_ > post-early_ign_, p < 0.001, 95% CI of the difference = 75.52 to 99.16) for the ignore condition, but not for post-early vs post-late timepoints (post-early_ign_ = post-late_ign_, p > 0.79, 95% CI of the difference = −12.69 to 5.15). The compensate vs ignore comparison was significant in the post-late timepoint (post-late_cmp_ > post-late_ign_, p < 0.001, 95% CI of the difference = 39.47 to 70.03), demonstrating that the introduced pitch perturbation directly influenced behaviour.

### 3.2 fMRI Findings

#### 3.2.1 Simple tone reproduction

Consistent with previous findings (Segado et al. 2018), a statistical conjunction of cello playing and singing compared to rest showed that both tasks recruited a bilateral network including M1 and dPMC, as well as SMA and STG (Figure 5).

**Figure 5.**
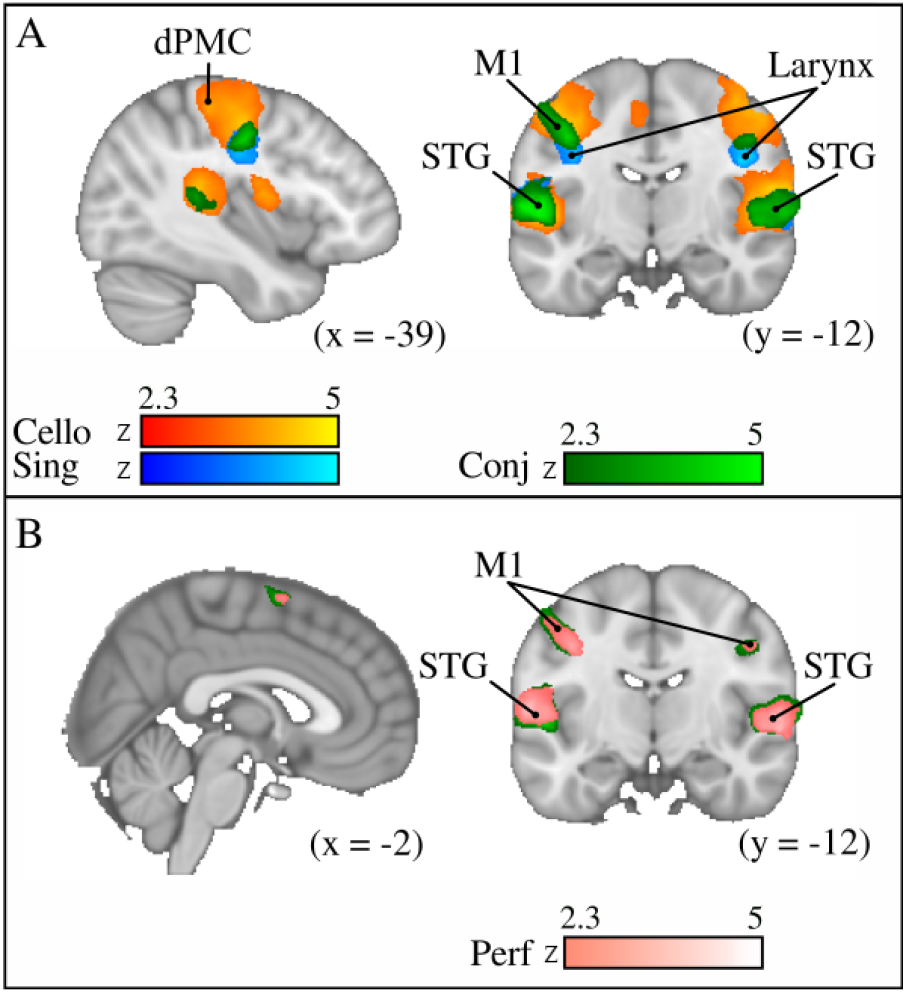
Simple performance condition | A) Simple cello playing (orange), singing (blue) and conjunction (green). B) Conjunction (green) and areas that were positively correlated with good task performance (pink). Singing and cello playing were found to have overlapping activity in pre- post-central gyrus, STG and SMA. This activity was found to be positively correlated with good performance on the task.

Additionally, singing specifically activated larynx area of M1, whereas cello playing recruited a region of M1 related to hand and arm movements. Activity in larynx area of M1 was identified based on bilateral 10 voxel spheres centered on the larynx area defined in Brown et al. 2006. Hand area was identified based on bilateral 10 voxel spheres centered on hand knobs (Yousry et al. 1997).

#### 3.2.2 Compensation

Only trials where participants successfully compensated for the perturbation were included in this analysis. Successful compensation was defined as returning to within 50 cents of the target tone by the end of a given trial. When comparing the Shift to the Simple no shift condition both singing and playing recruited a bilateral network of regions in the dorsal-stream: dPMC, pre-central gyrus, IPS, SMG extending to pSTG, and SMA extending to pre-SMA. These regions all correspond to those of the auditory-vocal integration network reported previously for the equivalent manipulation carried out in trained singers (Zarate et al. 2008). A statistical conjunction of Sing and Play for the Compensate condition showed that similar regions of SMA/Pre-SMA, dPMC, IPS, SMG were significantly active for both production modalities (Figure 6A). Regression analysis examining the relationship between compensation accuracy and BOLD signal from the Compensate vs Simple contrast in the regions of overlap showed that greater activation in all of these regions was positively correlated with pitch accuracy (Figure 6B).

**Figure 6.**
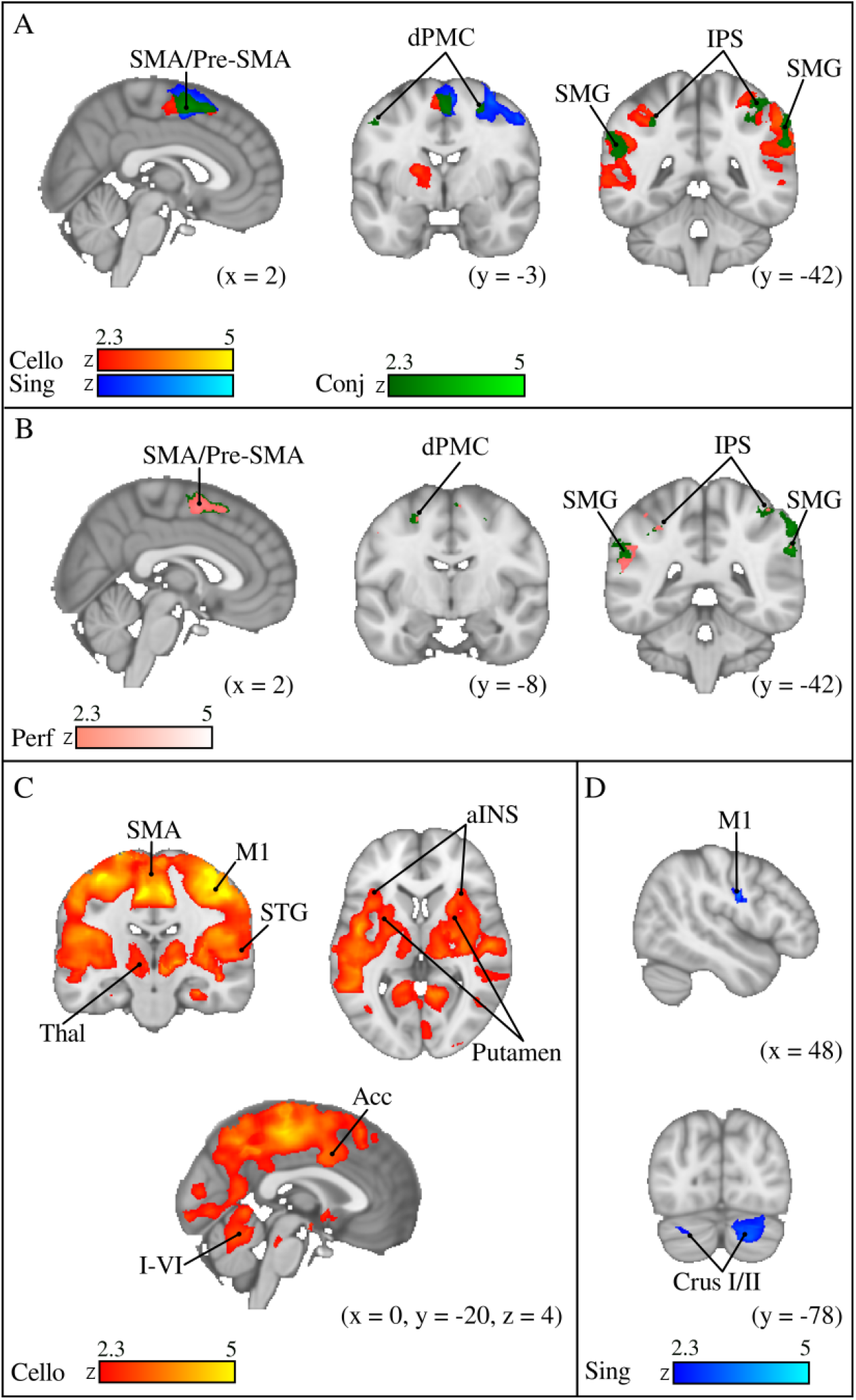
Compensating for pitch perturbations | A) Cello playing (orange), singing (blue), and Conjunction (green) of Compensate vs Simple contrast B) Areas within this conjunction (green) where activity is positively correlated with good task performance (pink) C) Cello playing > singing and D) singing > cello playing. Conjunction and regression show that SMA/pre-SMA, dPMC, SMG, and IPS are all contributing to good task performance in singing and in cello playing. Cello playing recruits more activity throughout the auditory motor integration network including in the BG and aINS. Singing shows more activity in vocal areas of motor cortex and cerebellum.

There were also significant differences observed between singing and cello playing in the Compensate vs Simple contrast. Cello playing showed stronger, and more extensive activation in bilateral dorsal motor and pre-motor regions extending posteriorly into the superior parietal lobule as well as in the SMA, parietal operculum, the pre-cuneus and bilateral cerebellar Crus V (Figure 6C). Singing preferentially recruited larynx area of M1, as well as cerebellar Crus I and Crus II (Figure 6D).

#### 3.2.3 Ignoring: similarities and differences

In the Ignore condition, singing and cello playing both recruited dPMC and STG when contrasted with Rest (Figure 7A). The statistical conjunction of Sing and Play for the Ignore vs Rest contrast showed overlapping activation in pSTG and dPMC. Regression analysis examining the relationship between accuracy and BOLD signal from the Ignore vs Simple contrast in the regions of overlap showed that greater activation in pSTG was positively correlated with performance score (Figure 7B).

**Figure 7.**
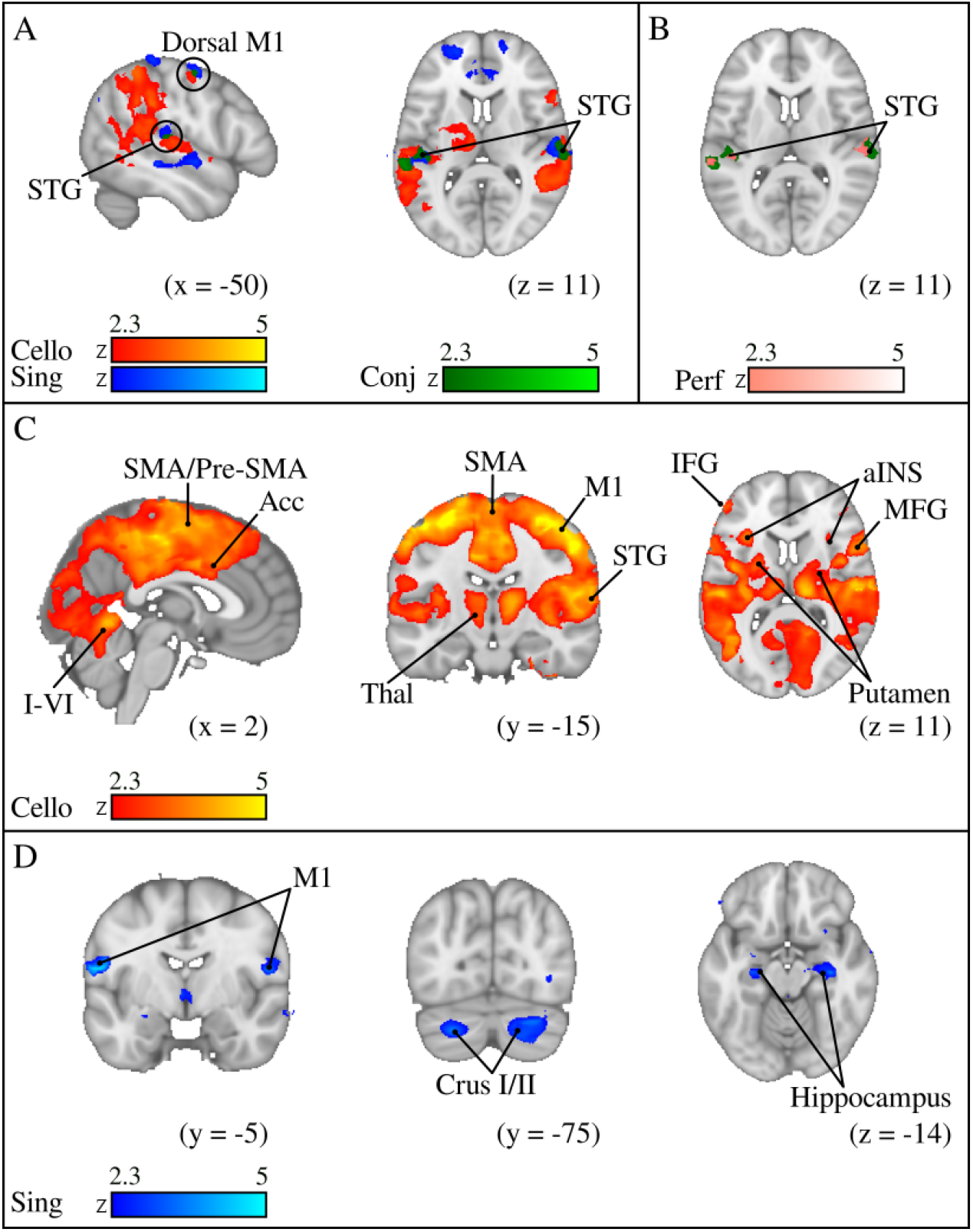
Ignoring pitch feedback perturbations | A) Cello playing (orange), singing (blue), and Conjunction (green) of Ignore vs Simple contrast B) Areas within this conjunction (green) where activity is positively correlated with good task performance (pink) C) Cello playing > singing and D) singing > cello playing. Conjunction and regression show that only posterior STG is contributing to good task performance in both singing and in cello playing. Cello playing recruits more activity throughout the auditory motor integration network including in the BG and aINS, MFG, and IFG. Singing shows more activity in vocal areas of motor cortex and cerebellum, and in the hippocampus.

As in the Compensate condition, cello playing elicited greater and more widespread activation relative to singing throughout bilateral dPMC extending posteriorly to the superior parietal lobule, in the SMA, the parietal operculum, and the pre-cuneus as well as in bilateral Crus V, VIIIa and VIIIb of the cerebellum (Figure 7C). Singing showed greater activation in larynx area and in Crus I/II of the cerebellum. In addition, singing showed greater activation in bilateral hippocampus (Figure 7D).

#### 3.2.4 Functional Connectivity

Functional connectivity analyses were carried out for all three conditions to determine if brain activity in the region of primary auditory cortex [Heschl’s gyrus (HG)] was correlated with activity in motor regions (or elsewhere) and, in doing so, determine whether multiple brain regions are working together in order to accomplish the experimental task (Figure 8). For both Sing and Play in all three conditions (Simple, Compensate, Ignore), activity in the HG seed was correlated with activity in bilateral STG within and around the HG seed, as well as with activity in the SMA, and the dorsal part of M1. For singing, HG activity was correlated with activity in regions of M1 corresponding to vocal articulators. For cello playing, HG activity was correlated with activity in regions of M1 corresponding to hand knob (bilateral).

**Figure 8.**
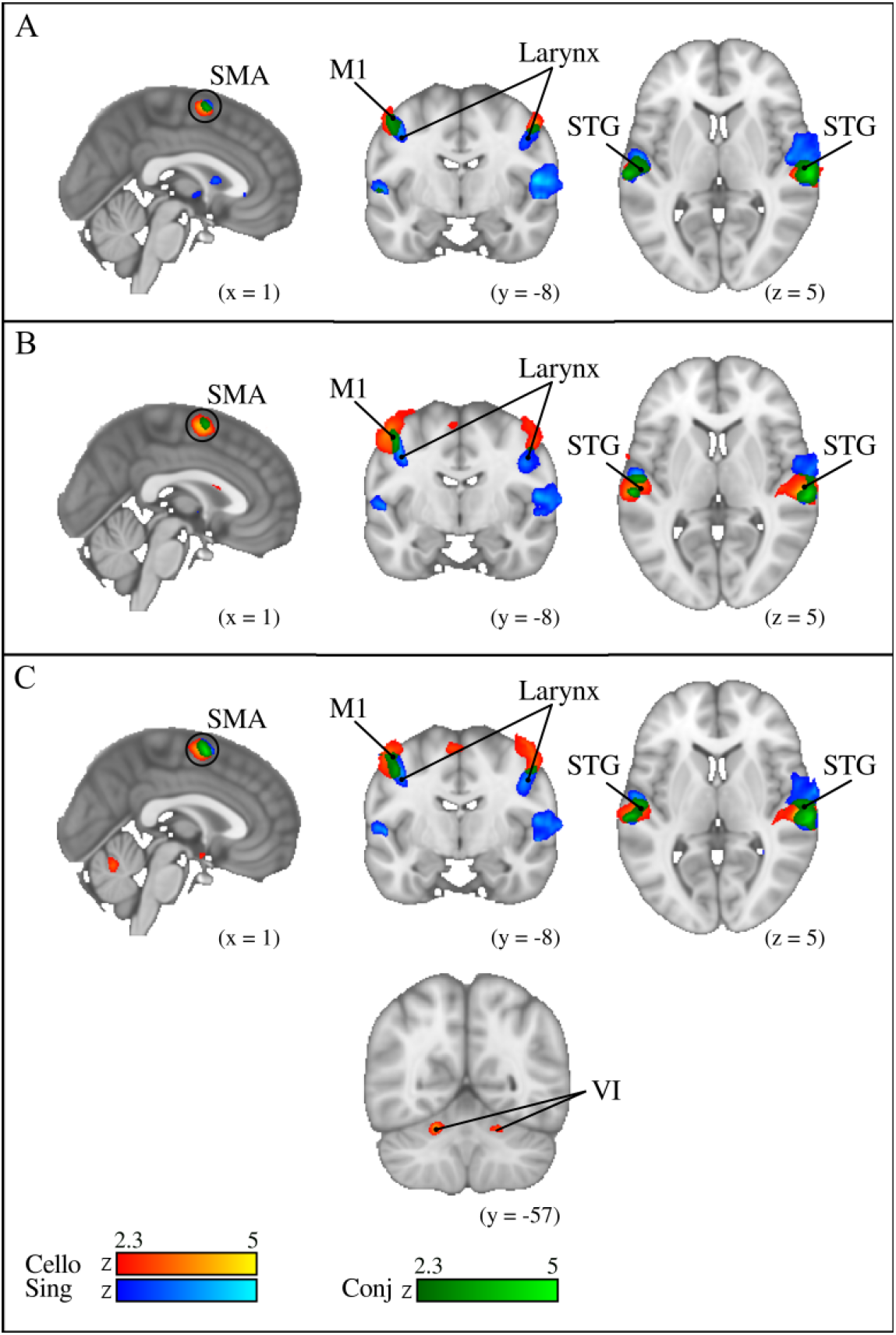
Functional connectivity from HG seed for cello playing (orange), singing (blue), and Conjunction (green) | A) Simple condition B) Ignore condition and C) Compensate condition. In all three conditions, activity in auditory cortex was correlated with activity in and around the HG seed as well as with activity in SMA and motor cortex. Activity in motor cortex was effector-specific with singing recruiting larynx area and cello playing recruiting hand area (both bilateral). In the compensate condition, only cello playing showed functional connectivity from HG to cerebellum.

## 4 Discussion

The findings of this study support our main hypothesis that similar brain networks underlie auditory-motor integration for singing and cello playing. First, the participants, who were expert cellists but not expert singers, were all able to accurately match target tones in the simple play or sing conditions and to accurately compensate or ignore auditory feedback perturbations for both production modalities. Second, the network of brain regions recruited for simple singing and cello playing directly overlapped, except in primary motor cortex where activity was observed in the larynx area for singing and the hand area for cello playing. Within the regions of overlap, BOLD signal change in auditory cortex and functional connectivity from an HG seed to a subset of these regions (STG, SMA, M1) were both positively correlated with pitch accuracy, indicating that the synchronous activity of this network of brain regions positively contributed to performance. Third, compensating for pitch perturbations when either singing or playing recruited a bilateral dorsal network (dPMC, IPS, SMG, pSTG, SMA/pre-SMA). BOLD signal change within all of these dorsal-stream regions was positively correlated with pitch compensation performance. Further, activity in HG was temporally correlated with activity in a subset of these regions (SMA, M1, STG) and bilateral Lobule VI of the cerebellum. Finally, when ignoring pitch perturbations for both singing and cello playing participants recruited overlapping regions in dPMC and pSTG. BOLD signal change and functional connectivity from an HG seed to pSTG was positively correlated with pitch accuracy score. Taken together these results indicate that there is a common network underlying auditory-motor integration that spans vocal-motor and hand motor control. This conclusion is consistent with the neuronal recycling hypothesis which posits that phylogenetically newer abilities dependent on human culture are scaffolded on phylogenetically older capacities (Dehaene 2005).

### 4.1 Overlapping brain networks for singing and playing

Our first finding, that performance was highly accurate for singing and cello playing across all three conditions (simple, compensate, ignore), is the first direct evidence showing that the magnitude of the behavioural response to pitch feedback perturbations in continuous pitch instruments is directly comparable to the response observed in singing. This finding is critical for interpreting our neuroimaging findings, as it allows us to more confidently attribute the observed similarities/differences in brain activity for these two tasks to specific brain networks, and not to underlying behavioural differences.

In the simple play/sing condition, the contrasts of singing and cello playing compared to rest, together with the conjunction analysis showed overlapping recruitment of brain areas throughout dorsal motor regions and the STG, with overall greater activation for playing than singing. We also observed expected differences in the location of motor activity, with singing activating the larynx area to a greater extent than playing, and playing activating hand-motor areas to a greater extent than singing. This is consistent with our previously published neuroimaging finding showing substantial overlap in the auditory-motor network for single tone reproduction when singing and playing (Segado et al. 2018). Unlike in that prior study, however, we did not see any significant activation in basal ganglia or brainstem despite their known role in pitch regulation and vocalization (Zarate et al. 2008). We attribute this to the use of a sparse sampling design that was not optimized to catch the peak of the hemodynamic response, meaning that signals that already had lower signal strength to begin with, like in the basal ganglia and brainstem, likely did not reach significance.

The compensate and ignore conditions give complementary information related to the feedforward and feedback components of the auditory-motor integration network, with the compensate condition weighted towards regions that are involved in feedback integration. The finding that cellists compensate as accurately when singing as they do on the cello is in line with research showing that non-singers compensate as accurately for large pitch shifts as do expert singers (Zarate et al. 2008), and demonstrates that auditory feedback is sufficient for pitch regulation in both tasks even when other sensory modalities (kinesthetic, vibrotactile) provide conflicting information. For the compensate condition we found directly overlapping brain activation in the IPS and SMG (part of the IPL) during singing and playing. Research has shown that the IPS and IPL (SMG) are recruited for pitch regulation tasks across a broad range of contexts including in singing (Zarate et al. 2008), speech vocalizations (Toyamura 2007), and piano performance (Pfordresher 2014, Brown 2012). A more global role for the IPS in pitch-related transformations is demonstrated by studies showing that it is engaged during performance of a transposed melody discrimination task, and that the active region overlaps with that involved in temporal re-ordering of pitches (Foster 2010, Foster 2013). It is therefore reasonable to speculate that for tasks requiring very similar computations, in this case the transformation of pitch feedback into a feed-forward motor command, similar infrastructure within the IPS would be used.

In the present study we show that, within the IPS and IPL, cello playing and singing recruit directly overlapping regions for the compensate condition. Within the IPS, overlap was seen primarily in anterior portions of the IPS, which have been found to relate to sensorimotor transformations and motor execution (Culham and Valyear 2006). This overlapping activity extended ventrally to the IPL which, as mentioned above is consistently found to be active in the musical instrument literature (Bangert 2006) at least in part due to its role in skilled object-related actions like tool use (Binkofski, Klann, and Caspers 2016; Culham and Valyear 2006). The mid to anterior localization of activity within the IPL is consistent with work showing that these regions are most related to motor planning and execution, simple motor behaviours, tactile reception, and a number of speech tasks like phonological short term memory (Tremblay, Shiller, and Ostry 2003). Given this, it is reasonable to interpret our finding as evidence that vocal and non-vocal tasks requiring pitch transformations for the purpose of movement planning, perhaps, have similar underlying computations and are therefore carried out within the same region largely independently of the source of auditory feedback and target motor effector. It is nonetheless possible that the current paradigm and resolution of the scanning protocol was not adequate to detect subtle differences in spatial locations of active regions, and that brain regions are tightly adjacent as opposed to directly overlapping. However, that would still suggest an organization within the IPS based on higher-level sensory transformations as opposed to motor effector. This question could be addressed directly in the future using a multivariate analysis technique like multivoxel pattern analysis that is more sensitive to differences between tasks within a region of interest.

The findings from the ignore condition allow us to further determine the extent to which singing and playing engage similar brain regions when auditory feedback is incorrect and performance is, as a result, based on a forward model. We found that participants performed the ignore task with as high a degree of accuracy when singing (for which they had no specific expertise) as they did when playing (for which they had a very high level of expertise). This finding suggests that the ability to successfully rely on a forward model is at least partly independent of the effector used to accomplish a sensory goal, and may be more related to feedback monitoring strategies than forward motor planning. We found that activity associated with ignoring in the two production modalities overlapped in dPMC and in pSTG, although there were also separable activations in the motor hand area and the larynx area for playing and singing respectively. Both dPMC and pSTG are consistently active in error monitoring and pitch control tasks (Baumann 2005, S. Brown 2004), with dPMC being specifically relevant for associating sounds with movements (Brown 2006) and pSTG being involved in error correction (Pfordresher and Mantell 2014, Tourville and Guenther 2011). We found that activity in pSTG was positively correlated with good performance, but activity in dPMC was not. Our interpretation of this correlation is that the increased activity in auditory cortex resulted from the absence of the typical inhibition of STG typically seen for monitoring of self-generated sounds (Bendixen, SanMiguel, and Schröger 2012; Sanmiguel, Todd, and Schröger 2013; Mathias, Gehring, and Palmer 2019; Christoffels, Formisano, and Schiller 2007). If participants were no longer linking the perceived sounds to their produced output, and were instead increasing their reliance on a forward model and on non-auditory sensory feedback, then it would follow that inhibition of auditory cortex would be weaker and activity in pSTG would be stronger as a result. We would therefore propose that the mechanisms involved in representing the target pitch stably in the face of competing inputs, and perhaps inhibiting corrective responses, are shared for both singing and cello, just as the mechanisms for compensation also appear to be shared.

### 4.2 Significance for feedforward and feedback control models

The extensive overlap observed for singing and cello playing across experimental conditions suggests shared mechanisms that can be used flexibly for pitch regulation. One interpretation is that, consistent with the theory of neuronal recycling, these mechanisms originally developed for vocal control and are now being used for the purpose of pitch regulation during instrument playing. The vocal pitch regulatory system, including several auditory-vocal reflexes, are present in phylogenetically older species like non-human primates (Eliades and Wang 2012), bats and frogs (J. Luo, Hage, and Moss 2018). Given that none of these species play musical instruments (yet), they would have few if any uses for pitch regulation outside the context of vocalization and it therefore seems reasonable to speculate that the brain mechanisms governing vocal pitch regulation developed primarily for that purpose. However, a complementary interpretation is that both vocal control and instrument playing make use of more general-purpose feedforward and feedback mechanisms that are important for the control and regulation of movement. Research done on visuomotor control has shown that error correction may be best modelled as a Bayesian process (Kording and Wolpert 2004) with the strength of the sensory prior, along with incoming sensory information, determining whether or not a corrective action takes place. Such a framework has been shown to account the observation that expert singers are better able to ignore altered pitch feedback while still showing a reflexive compensatory response to small pitch shifts (Zarate 2010; Hahnloser and Narula 2017).

## 5 Conclusion

We found that cellists can perform pitch perturbation tasks as accurately when playing the cello as when singing, and that playing/singing make use of directly overlapping brain areas both when compensating for and ignoring introduced pitch perturbations. The functionally connected network of overlapping brain regions includes those used for auditory feedback processing and auditory-motor integration, but differs at the level of forward motor control. This finding suggests that regions responsible for auditory-motor integration, like the IPS and SMG, are performing high level sensory transformations as opposed to coding for specific motor actions. Moreover, we found that the network of overlapping brain regions is consistent with the singing network described in the literature, and with work done in auditory-motor integration during musical instrument performance, further suggesting that brain networks for the two tasks may be shared. We propose that this is due to a co-opting of brain mechanisms that developed primarily for the purpose of vocalization, which is consistent with the neuronal recycling hypothesis.

## 1. Author Contributions

The authors all contributed to the conception and design of the study. Melanie Segado collected and analysed both the fMRI and behavioral data, including statistical analyses, and wrote the first draft of the manuscript. All authors contributed to manuscript revision, read and approved the submitted version.

## 2. Conflict of Interest Statement

The authors declare that the research was conducted in the absence of any commercial or financial relationships that could be construed as a potential conflict of interest.

## 3. Acknowledgments

The authors gratefully acknowledge the work of Yilin Zhang and Vanessa Mukli for their help recruiting and scheduling participants, collecting data, and with data cleaning and analysis. They further acknowledge the work of Joseph Thibodeau for software development and technical support. This work was supported by an operating grant from the Canadian Institutes of Health Research to Robert J Zatorre and Virginia B Penhune and by an infrastructure grant from the Canada Foundation for Innovation. Robert J. Zatorre is a fellow of the Canadian Institute for Advanced Research.

## Notes

### Competing Interest Statement

The authors have declared no competing interest.

